# Dynamic adaptation to novelty in the brain is related to arousal and intelligence

**DOI:** 10.1101/2024.08.02.606380

**Authors:** Jacob Tanner, Joshua Faskowitz, Daniel P. Kennedy, Richard F. Betzel

## Abstract

How does the human brain respond to novelty? Here, we address this question using fMRI data wherein human participants watch the same movie scene four times. On the first viewing, this movie scene is novel, and on later viewings it is not. We find that brain activity is lower-dimensional in response to novelty. At a finer scale, we find that this reduction in the dimensionality of brain activity is the result of increased coupling in specific brain systems, most specifically within and between the control and dorsal attention systems. Additionally, we found that novelty induced an increase in between-subject synchronization of brain activity in the same brain systems. We also find evidence that adaptation to novelty, herein operationalized as the difference between baseline coupling and novelty-response coupling, is related to fluid intelligence. Finally, using separately collected out-of-sample data, we find that the above results may be linked to psychological arousal.

## INTRODUCTION

Adaptation to novelty is fundamental to life [1]. We continually encounter novel situations and information, and our ability to learn, adjust our behavior, and even reshape our mental models is crucial for our success and well-being. The ability to adapt to novel environments, respond to novel threats, and recognize novel opportunities has been central to the survival and diversification of species. This principle extends to the lives of individual organisms as well. For example, when an animal encounters a novel type of food or a novel potential predator, its survival may depend on its ability to adapt to this novelty. In humans, this ability to adapt is not only a biological process but also a psychological one.

The human brain is fundamentally a complex adaptive system. In complex systems, the local interactions of relatively simple parts result in complex and emergently adaptive behavior. In humans this feature allows us not only to regulate our homeostatic needs but also to build models of novel environments. Network science provides us with tools well suited for analyzing the interactions in such systems. By modeling the brain as a network, we can consider each element in this complex system as a node in a network and their interaction as defining the strength of the connection between them. Using the tools of network science, network neuroscience has shown that the brain is a hierarchically modular system where many units synchronize their activity to form communities at different scales: from cellular ensembles [2, 3], to brain regions, to brain systems [4–6].

The various scales in this hierarchically modular system define ways the system is often approximately lower dimensional. Previous work has described how such lower dimensional dynamics relate to the elements of tasks [7–9], and the building of environment-specific models [10]. Some work suggests that the brain changes dimensionality in the presence of stimuli [11]. Likewise, the modular structure of functional brain networks has been related to areal specialization of function [12, 13]. Furthermore, dynamic changes to this modular structure have been related to learning [14], task complexity [15], cognitive flexibility [16], surprise [17], and intelligence [18].

In this paper, we explore how modular human brain networks respond and adapt to novelty. As complex adaptive systems, our brains form and reform models of the environment in order to become more adaptive [19, 20]. In this way, novel stimuli could initiate a model learning process that helps to regulate the systems behavior. In this paper we model novel stimuli using a paradigm in which human subjects watch a movie scene on multiple occasions. The first time that human participants watch the movie scene is novel, while subsequent viewings are not. To the degree that participants are able to model this novel stimuli, we expect that brain dynamics reflecting this process should be different on the first viewing than on subsequent viewings.

We show that the first viewing of this movie scene initiates dynamic coupling between various brain systems that lowers the overall dimensionality of the system. We find that this decreased dimensionality occurs because brain systems like the control and default mode network temporarily increase their connectivity with other brain systems (like dorsal attention and visual systems) during the presence of novel stimuli.

We argue that these brain systems (e.g. control, and default mode) are assisting the brain in adapting to the novel stimuli by building model(s) of it. Concordantly, when the stimuli is no longer novel the activity of these brain systems becomes more independent from the stimuli, resulting in an increase in the dimensionality. Accordingly, we also show that this process of dynamic adaptation to novelty in the brain might be related to fluid intelligence. That is, an individual’s fluid intelligence is related to how much that individual adapts to and models novel stimuli.

Finally, we collected data from a new set of subjects that dynamically rated their level of arousal with a joystick while watching these movie scenes (outside of the scanner). We found that the novel viewing of the movie scene resulted in increased levels of psychological arousal, and an increase in across subject synchronization of dynamic arousal to the movie scene. Additionally, this dynamic synchronization of arousal was highly related to the out-of-sample activity in the thalamus, echoing previous work relating thalamic activity and arousal to the low-dimensional architecture of neural activity [15, 21].

Understanding how the human brain dynamically responds and adapts to novel information could be a fundamental first step to understanding how we build models of the world. Here we show that this process likely involves increased coupling within and between the control system, default mode network, and dorsal attention system alongside an increased between-subject synchronization of activity in a similar set of regions. Furthermore, we provide evidence that this response might be mediated by arousal systems involving the thalamus.

## RESULTS

In our study we analyzed data from 7 Tesla fMRI movie-watching data collected as part of the Human Connectome Project [22]. In these data, participants watched a selection of movie scenes (from both Hollywood movies and independent films) on four separate occasions. The final movie scene on each occasion was the same. Here, we analyze time series data from 129 of these subjects who watched this same movie scene four times (for a schematic describing the organization of these data, see Fig. 1a,b).

**FIG. 1.**
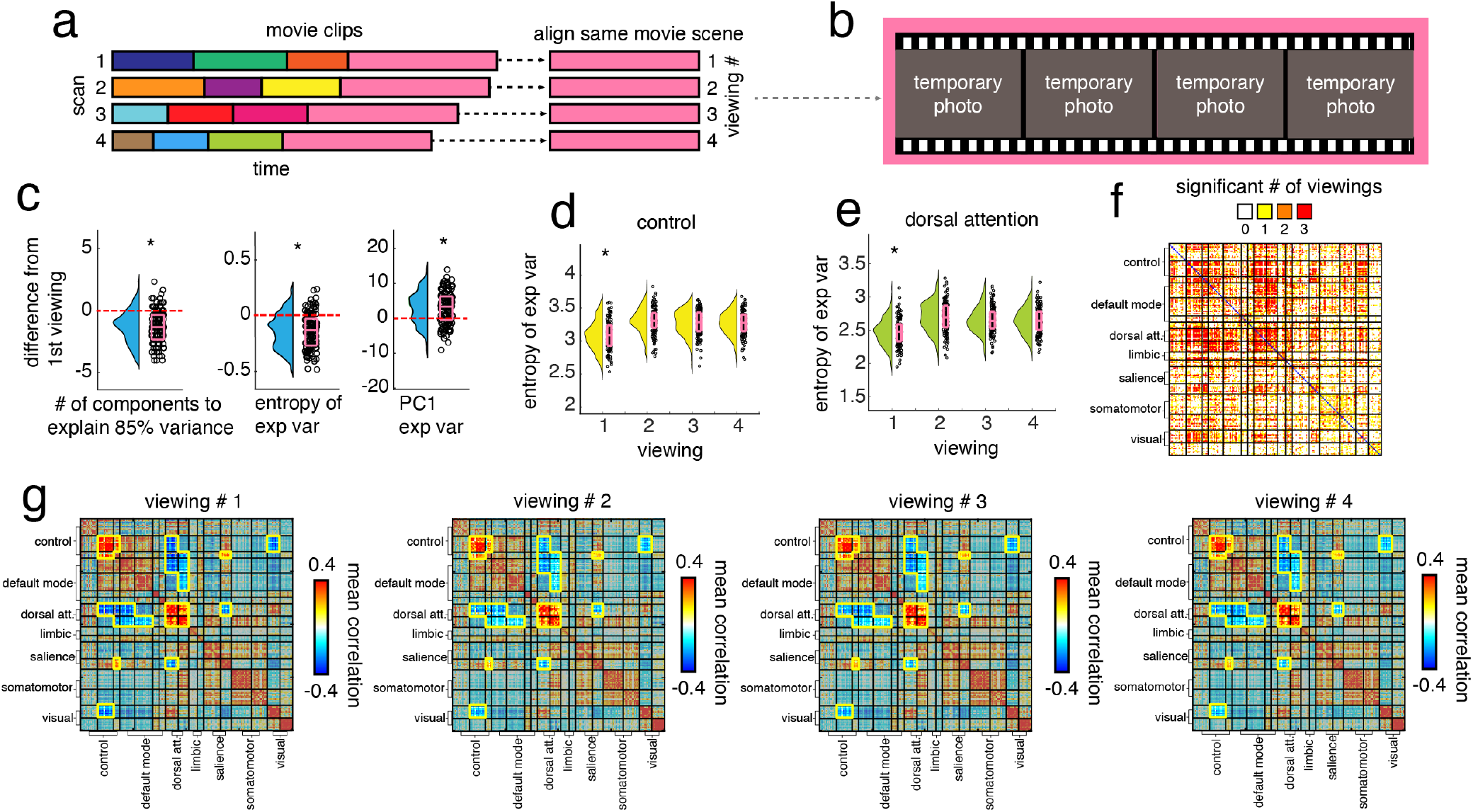
Novelty effects, (*a*) Schematic illustrating the structure of the fMRI data. Participants came in for four separate scans. During each scan, they watched a series of movie scenes. However, at the end of each scan was the same movie scene. We aligned the fMRI data from each of these viewings of this movie scene for analysis in the rest of the paper. (*b*) Sample of images from this movie scene. (*c*) Box plots showing the difference between (1) the number of principal components needed to explain 85% of the variance in brain activity, (2) the entropy of the explained variance distribution, and (3) the explained variance of principal component 1 during the first viewing and subsequent viewings (on average). (*d*) Box plots showing the entropy of the explained variance distribution of the control systems activity for all viewings. (*e*) Same but for the dorsal attention system. (*f*) Matrix showing the number of subsequent scans for which each edge had greater connectivity during the novel viewing. (*g*) Functional connectivity (mean across subjects) matrices for all viewings. System-by-system blocks with a significant number of novelty effects are outlined in yellow.

In this section we describe our evidence for the effect of novelty on the brain. We analyze data wherein human participants watch the same movie scene on four separate occasions, and show that the human brain responds differently on the first viewing of the movie scene. One common response to this novelty is to increase coupling between a select set of brain regions, effectively reducing the dimensionality of the system. Additionally, we find that the same brain systems that increase their coupling also increase their across-subject synchronization during the novel viewing.

### Decreased dimensionality and increased connectivity in response to novelty

In this section we show that the novel movie viewing tends to be related to decreased dimensionality and increased connectivity for a select set of brain systems. Dimensionality can be approximated by linear methods using principal component analysis (PCA) to investigate the distribution of explained variance. If the explained variance is mainly concentrated in a small number of dimensions, then the system is lower dimensional. We use three measures to explore this: (1) the number of components that it takes to explain 85% of the variance, (2) the entropy of the explained variance distribution, and (3) the explained variance of the first principal component.

We found that the first viewing of a novel movie scene resulted in decreased dimensionality by all three metrics (when compared to other viewings; Fig. 1c; *p <* 0.05).

Next, we performed PCA separately on each of seven brain systems defined by Schaefer et al [5] and found that the change in overall dimensionality was primarily due to changes in the dimensionality of the control system and the dorsal attention system (Fig. 1d-e; *p <* 0.05).

To investigate these effects in greater detail, we used functional connectivity analysis. Intuitively, this descreased dimensionality should result in increased connectivity during the first viewing. We therefore calculated the functional connectivity for every viewing, and compared the connectivity values of each edge for the different viewings. This involved collecting the edge strength for an edge across all subjects, and comparing this distribution on the first viewing with the distributions of later viewings. For a novelty effect to be consistent, this connectivity distribution should be higher (lower) on the first viewing than all three subsequent viewings.

For every edge, we calculated how consistent this connectivity effect was by taking the number of subsequent viewings the connectivity distribution was less than (greater than) the distribution in the first viewing (Fig. 1f; *p <* 0.05). We then used a spin test to identify brain systems (and their interactions) that had a significantly consistent connectivity-based novelty effect compared to a spatially constrained null model, and we found that control system B, dorsal attention systems A and B, as well as edges between control system C and the salience system B increased their connectivity on the first novel viewing (Fig. 1g, *p <* 0.05). In contrast, edges between the control system and the dorsal attention system, as well as between the control system and the central visual system, and salience system B and the dorsal attention system A had higher negative correlations on the first novel viewing (Fig. 1g, *p <* 0.05).

In order to ensure that these effects were not due to confounds, we tested a number of alternative possibilities. We found that the first scan of the resting-state scans showed no significant decreases in dimensionality (Fig. S2d, *p <* 0.05). Additionally, we also checked to see if subjects exhibited more motion on the first scan, and they did not (Fig. S1a-d). Further, we found that this novelty effect remained intact across different parcellation scales (200 and 400 nodes; Fig. S2a,b, *p <* 10^−5^). We also checked if this effect was still intact when no global signal regression is used during processing. We found that the first viewing had significantly lower dimensionality than the next two viewings, but not the final viewing (Fig. S2c, *p <* 0.05).

### Increased inter-subject synchronization in response to novelty

In this section, we show that the brain activity of participants watching the novel movie scene is more synchronized on the first viewing than later viewings. Additionally, the brain systems that were more synchronized were also those that we found increased their connectivity in the previous section.

In order to explore this, we first performed a within-subject analysis to identify which brain regions/systems had a similar temporal activity profile each time a subject watched the movie scene. We found that visual and dorsal attention systems tended to maintain similar temporal activity profiles each time that subjects watched the movie, but other brain systems did not (Fig. 2a-c, *p <* 0.05).

**FIG. 2.**
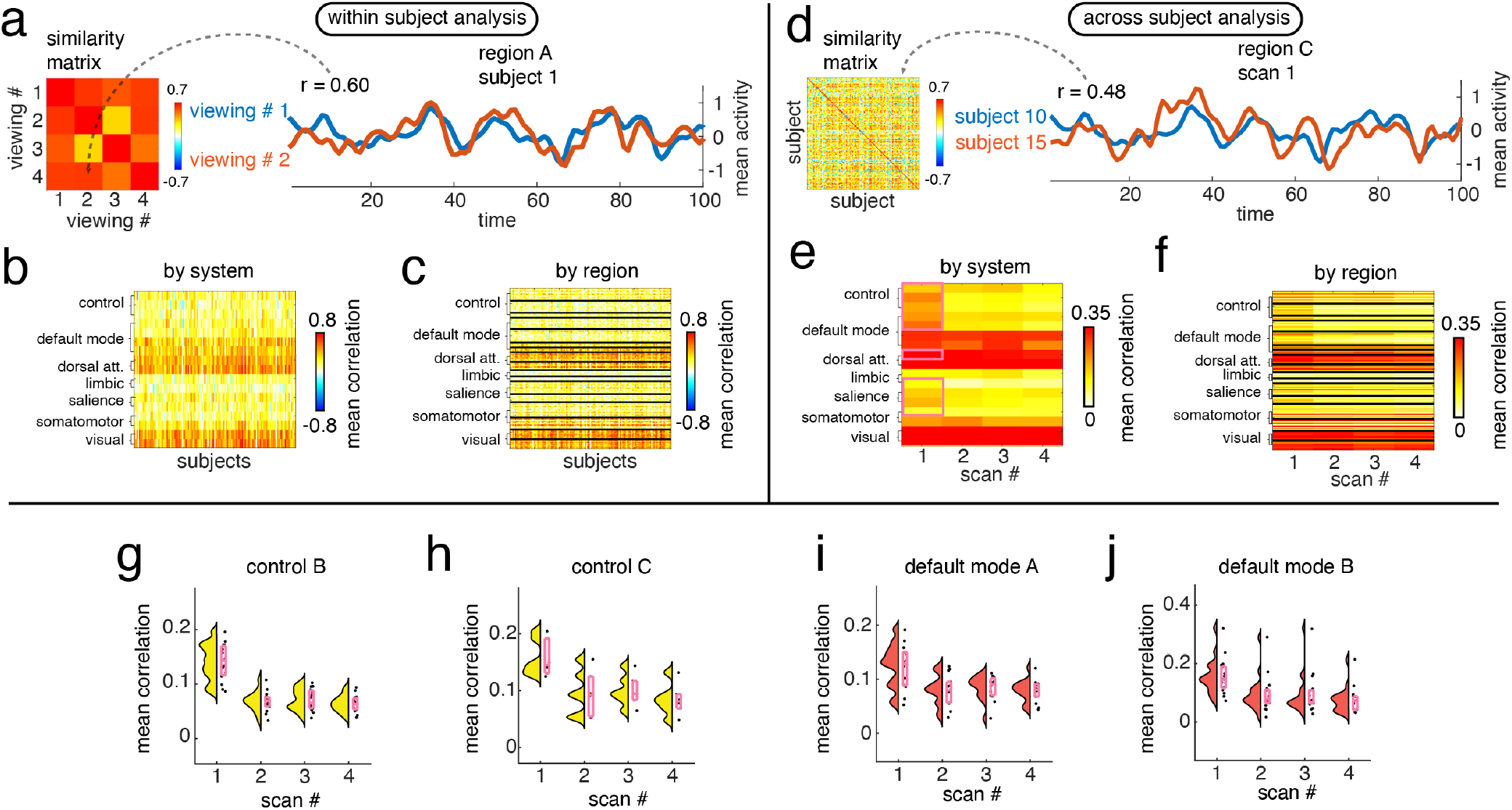
Increased inter-subject synchronization in response to novelty, (*a*) Schematic describing our within subject analysis. (*b*) Matrix showing the mean within subject (across viewing) correlation values for each system, and region (*c*). (*d*) Schematic describing our across subject analysis. (*e*) Matrix showing the mean across subject (same viewing) correlation values for each system, and region (*f*). Finally, we show box plots of the mean across subject correlation per scan/viewing for the control system B (*g*), control system C (*h*), default mode network A (*i*) and default mode network B (*j*).

Next, we performed an across-subject analysis to see if brain regions/systems were more or less synchronized across subjects on different viewings. We found that a large number of brain systems increased their across subject synchronization on the first viewing, including the control system (A,B,C), the default mode network (A,B), the dorsal attention system A, limbic system B, salience system (A,B) and somatomotor system A (Fig. 2d-f, *p <* 0.05), but the systems where this effect was the greatest were control (B,C) and default mode network (A,B) (Fig. 2g-j, *p <* 0.05).

In order to check if these results were due to a similarity in participants’ motion responses to the movie, we took the similarity between all subjects time-varying motion for each viewing. Each distribution per viewing represents how aligned motion dynamics are across subjects. All of these distributions were consistent with chance (Fig. S1).

Finally, in order to directly explore the similarity between the set of brain regions that increased their coupling (Fig. 1g-j) and the set of brain regions that increased across subject synchronization we performed an additional analysis wherein we created an inter-subject functional connectivity matrix.

Briefly, this matrix was created by computing the correlation between region *i*’s activity from subject *A* and region *j*’s activity from subject *B*, for all participants and all pairs of regions, *i* and *j*. We then calculated the mean value of all of these correlations to be the value of *edge*(*i, j*) in the inter-subject functional connectivity matrix. This matrix measures the inter-subject synchronization between all pairs of brain regions. We created one of these matrices for each viewing (Fig. 3c) and then compared the distribution of edges within and between each brain system.

**FIG. 3.**
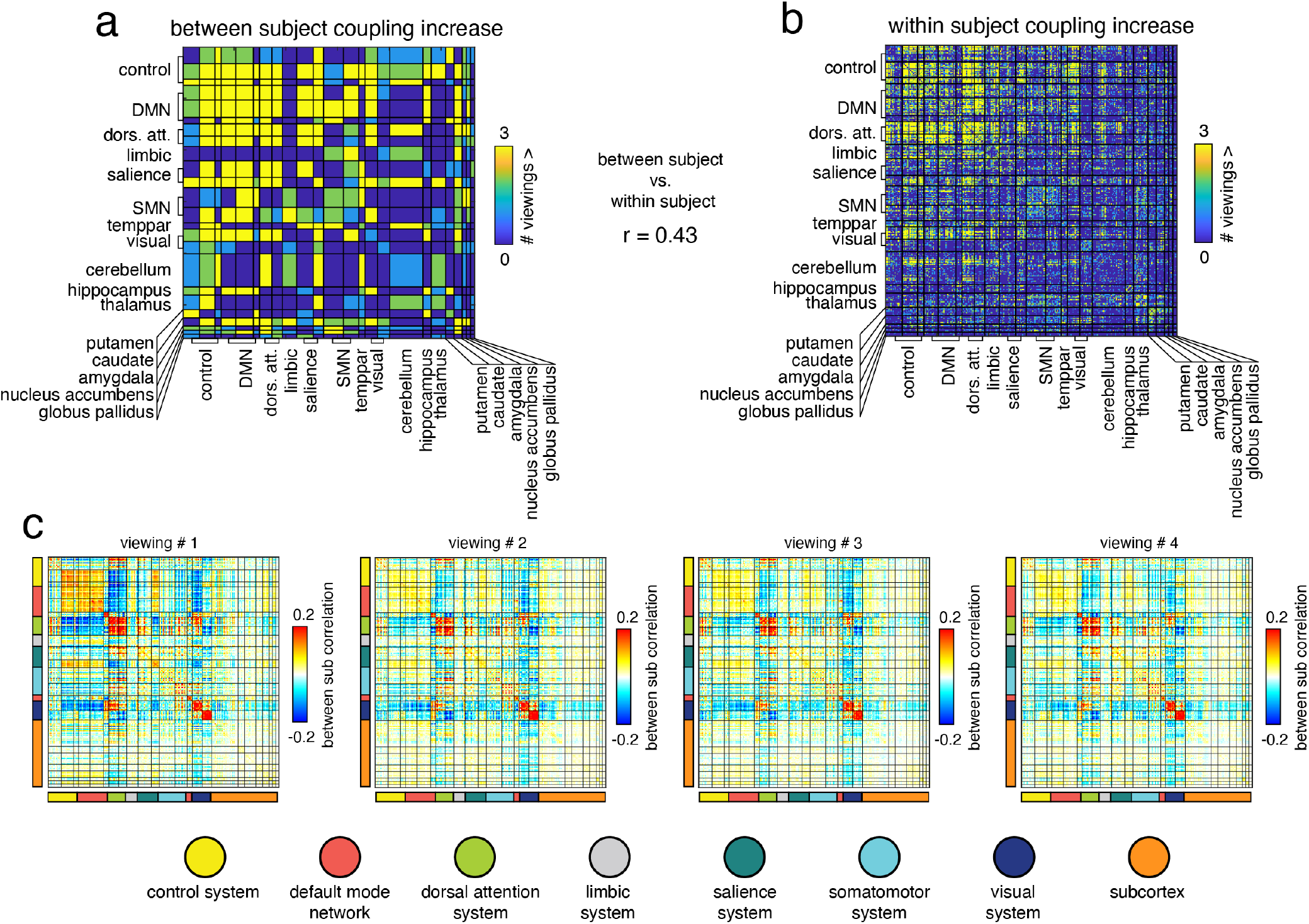
Between subjects versus within subject coupling, (*a*) Matrix showing, for each system-by-system block, the number of subsequent scans where between subject functional connectivity values were greater for the first viewing. (*b*) Matrix showing, for each edge, the number of subsequent viewings that had less connectivity than the first viewing. (*c*) Inter-subject functional connectivity for each viewing.

After counting the number of subsequent viewings where within or between system correlation values were significantly greater (or less) than the novel movie viewing, we had a matrix showing the number of significant values for each within, and between-system block (Fig. 3a). In support of our suspicion that the regions with increased coupling were also increasing their between subject synchronization, we found that this matrix was positively correlated with a matrix representing the edges with increased connectivity in the first viewing (versus subsequent viewings; *r* = 0.43, *p <* 0.05; Fig. 3b).

Taken together, these results suggest that novel movie scenes result in an increase in across-subject synchronization, and this phenomenon is highly related to the phenomenon of increased connectivity during novel movie scenes.

### Adaptation to novelty and intelligence

We hypothesized that the degree of adaptation to novelty in the brain would be related to fluid intelligence. In this section, we develop a measure of adaptation using the novelty effects we described in the previous section and show that this measure of adaptation is positively related to fluid intelligence.

To define adaptation, we measured the difference in connectivity between the first viewing and subsequent viewings. More specifically, we took the absolute difference between connectivity on the first viewing and the mean connectivity of subsequent viewings. Intuitively, this measure captures how different the novelty effect is from the baseline effect of the movie scene.

We hypothesized that brain systems that we identified in the previous sections would be most likely to have adaptation effects related to intelligence. Two conspicuous brain systems were the control system and the dorsal attention system. The connectivity within these systems not only increased in response to novelty, but the activity between these systems also became more strongly negatively correlated during the novel viewing. To investigate this, we took the mean activity within the control system and the dorsal attention system for each viewing, and took the correlation between these brain systems per viewing. To measure adaptation, we took the absolute difference between these correlation values on the first viewing and on later viewings Fig. 4a). We found that these adaptation values were positively related to subjects’ fluid intelligence scores (Fig. 4b; Spearman’s *ρ* = 0.23, *p <* 0.05). When we extended this analysis to consider every pair of brain systems, we found that this effect was the only one to be both positive and significant (Fig. 4c).

**FIG. 4.**
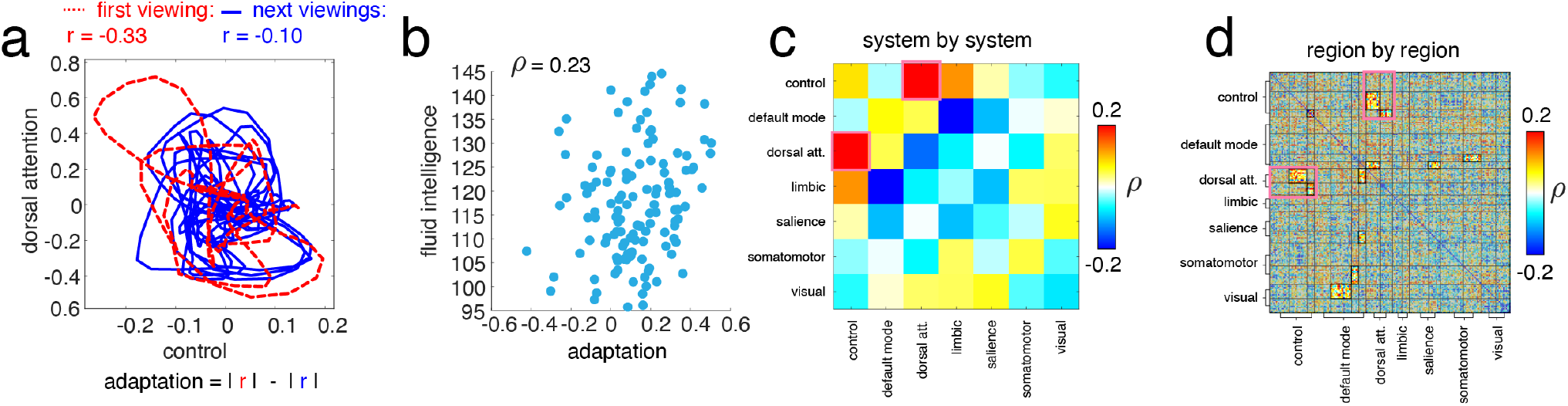
Adaptation to novelty effect in the brain is related to intelligence, (*a*) Plot showing the mean activity (across subjects) in the control system, plotted against the mean activity in the dorsal attention system. The trajectories are colored according to whether they happened in the first viewing (red) or on subsequent viewings (blue). (*b*) Plot showing the relationship between adaptation (absolute difference in connectivity between first and later viewings) and fluid intelligence. (*c*) Plot showing the correlation between adaptation and fluid intelligence for different pairs of brain systems, and regions (*d*). For the region by region correlation matrix, we conducted a spin test to assess if significant correlations were concentrated in any system-by-system blocks. Blocks that did not pass the spin test are greyed out.

Finally, we wanted to investigate if this adaptation effect and its relationship with fluid intelligence persisted at a finer scale involving the coupling of individual brain regions. We took the full functional connectivity network of each individual on the first viewing and on subsequent viewings and calculated the adaptation effect for each edge and subject. We then correlated these adaptation values with fluid intelligence and found that those correlations that were strong and significant tended to be concentrated in blocks connecting the control system to the dorsal attention system, as well as blocks connecting default mode network to visual, somatomotor, dorsal attention and salience networks (Fig. 4f,g; edges with *r >* 0.2 and *p <* 0.05 were tested using a spin test, *p <* 0.05), thus supporting that this adaptation effect persisted at a finer scale.

### Increased psychological arousal in response to novelty

We hypothesized that one factor that could partially drive novelty effects in the brain could be arousal (e.g. [23]). Perhaps systems that instantiate this effect in the brain modulate cortical coupling to produce the various effects described above. To explore this, we collected data from 46 subjects who watched all of the same movie scenes from the original fMRI dataset while dynamically rating their level of psychological arousal with a joystick. More specifically, the subjects watched all of the movie scenes described in Fig. 1a, including viewing the repeated movie scene four times.

We found that subjects reported more overall arousal on the first viewing of the novel movie scene (Fig. 5a; one-sample t-test; *p <* 0.05). Next, we tested and confirmed that these arousal ratings were more synchronized across subjects during the first viewing (Fig. 5b; one-sample t-test; *p <* 0.05).

**FIG. 5.**
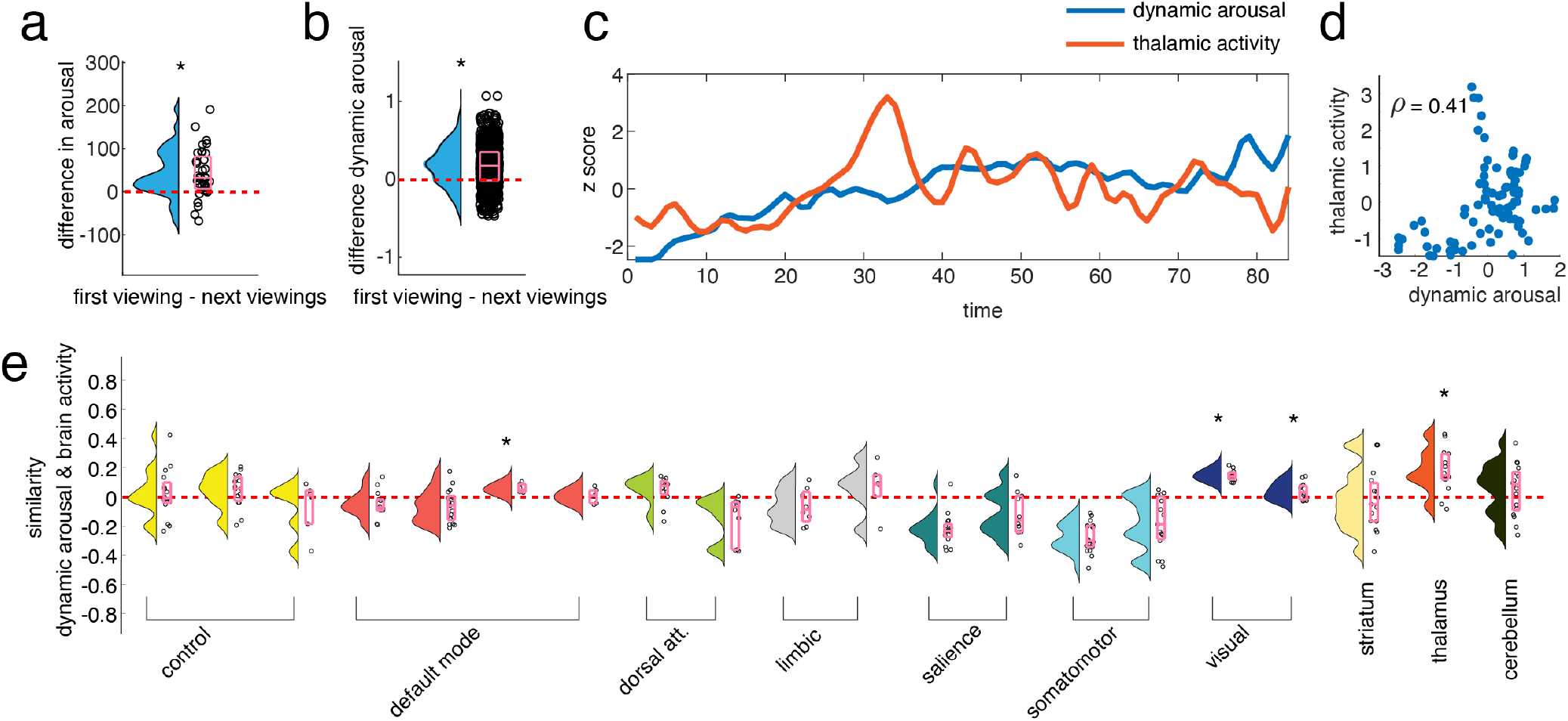
Adaptivity and arousal, (*a*) Box plot showing the difference between average arousal during the first viewing and average arousal on subsequent viewings. (*b*) Box plot showing the difference between inter-subject arousal synchronization (dynamic arousal) during the first viewing and subsequent viewings. (*c*) Plot showing the average arousal trajectory during the first viewing, and the average thalamic activity. (*d*) Plot showing the average arousal trajectory on the first viewing plotted against the average thalamic activity. (*e*) Box plots of the correlation of average arousal and activity in each region of different brain systems on the first viewing. Asterisks indicate systems whose distribution of correlation values were significantly greater than zero.

In order to investigate if a signature of this arousal effect could be found in the brain activity of subjects who watched these movie scenes in the scanner, we compared dynamic arousal ratings with brain activity. Given that previous research has suggested that that thalamus is involved in arousal [24–26], we chose to focus on the relationship between arousal ratings (outside of the fMRI scanner) and thalamic activity.

To do this, we took the mean of the dynamic arousal ratings across subjects in order to get an estimate of the average populations dynamic arousal response to watching this movie scene for the first time. Then, we correlated this average temporal profile of arousal with the average temporal thalamic activity profile across subjects who watched this movie in the scanner. We found a positive correlation between the mean activity of the thalamus and this mean measure of dynamic arousal (Fig. 5c; Spearman’s *ρ* = 0.41, *p <* 0.05). This effect was not present for later viewings.

In addition, we checked to see if this effect could be found in any other brain systems. To check this, we correlated the activity of every region (cortical and subcortical) with this mean measure of dynamic arousal on the first viewing. We found that the activity of regions in the default mode network C, both central and peripheral visual systems, and the activity of the thalamus were more related to dynamic arousal than chance (Fig. 5e, one-sample *t*-test, *p <* 0.05).

## DISCUSSION

How does the brain respond to novelty? Early research by Eugene Sokolov and others on the novelty response, or “orienting reflex” found that humans and other animals would involuntarily attend to novel stimuli in the environment until they gradually habituated to that stimuli, as he hypothesized, by developing neural representations to model the novel input [23, 27–32]. Later, it was found that the characteristic neural accompaniment to this orienting response was the novelty P3, an eventrelated brain potential (ERP) involving activation of the frontal and parietal cortices [23, 27, 33–35], although experiments combining ERP and fMRI also found effects in the superior temporal gyrus [36].

Neuroscience and psychology studies have also studied novelty processing generally as an important feature of cognition and adaptation [37–42]. For example, recent research in the primate brain found that novelty is intertwined with computations of sensory surprise and recency, also finding activation in temporal areas, specifically posterior medial temporal cortex [43]. Additionally, a 2015 review on novelty processing and its effects on the brain and cognition suggests that novelty increases arousal, improves perception and action, increases motivation and exploratory behavior, and promotes learning, also implicating a large variety of brain regions including the temporal and frontal cortices [44]. Additionally, modern innovations in machine learning have seen major improvements by integrating conceptions of novelty-seeking and detection into both agents and optimization processes [45–48].

Here, instead of focusing on individual novel stimuli (as many of these past studies have), we used more dynamic novel stimuli in order to characterize how the novelty response unfolds across time with more naturalistic stimuli. We operationalized this question using neuroimaging data from human participants repeatedly viewing a novel movie scene, a paradigm that has been used several times in fMRI literature (e.g. [49–53].

Our results suggest that this novel movie scene induces a decrease in the overall dimensionality of brain activity that involves in increased coupling within and between a select set of brain regions, primarily in the control and dorsal attention systems. These systems involve brain regions such at the inferior parietal lobule, lateral dorsal/oi9ventral prefrontal cortex, and medial posterior prefrontal cortex in the control system, and temporal occipital, parietal occipital, superior parietal lobule, post central, and frontal eye fields in the dorsal attention system. This result is in line with areas of interest involved in the orienting response, and also aligns with more recent results using repeated novel stimuli wherein the authors found that lateral temporal and inferior frontal regions cluster together more during the first viewing of novel stimuli [50]. Interestingly, we find that the neural trajectories that characterize this increased connectivity appear to synchronize across participants. This suggests that the novelty response is time-locked to the shared stimuli (the novel movie scene).

Following the first viewing of the movie scene (as the movie scene is no longer novel) later viewings tend to evoke somewhat different dynamics. The dimensionality of the activity increases, primarily in the control and dorsal attention systems, and a similar set of brain regions becomes less synchronized across subjects, suggesting these regions are either no longer tracking the stimuli or they are now tracking the stimuli in a more idiosyncratic manner. Nonetheless, more primary visual regions tend to maintain a similar set of dynamics for tracking the stimuli, and as such remain synchronized across subjects for all viewings.

Presumably, one of the primary functions of our brain is to form and reform models of the environment in order for an organism to become more adaptive. Many different theories about organisms and brains generally agree on the importance of modeling the environment, but disagree on the nature of the models and their function (e.g. [19, 20, 54–59]).

These results present a thought provoking picture of the neural response to novelty in human beings that might be unified by considering how they relate to the model building process. For example, some research suggests that the thalamus plays a key role in mediating the relationship between attention and arousal [26]. The presence of both increased arousal, and the link to thalamic activity in our results could suggest that participants are increasing their attention to the novel stimuli for the purpose of model building. Indeed, research on the orienting response in human beings suggests that we can only recognize novelty by comparing incoming sensory stimuli with a model, and that differences between the stimuli and the model provoke an increased attention to the novel stimuli for the purpose of model building [23].

Model building and increased attention might also explain why we see an increased between-subject synchronization in response to novel stimuli. That is, as participants pay more attention to the movie scene on first viewing for the purpose of model building, their brain activity becomes more time-locked to the movie. Likewise, on later viewings, the brain activity of participants (mostly in control systems) is no longer as time-locked to the stimuli because the model building process has mostly been completed.

This model building hypothesis also helps to explain the relationship between our measure of adaptation and fluid intelligence scores. This can be most clearly illustrated in reference to Raven’s Progressive Matrices [60–63], the test often used to measure fluid intelligence (a variant of this test was used to compute the scores used in this study, namely Penn Progressive Matrices [64]). Raven’s Progressive Matrices presents subjects with novel sequences of designs following a specific pattern. The task of the participant is to model the sequence and to use the inferred pattern to complete a partial sequence.

In this way, fluid intelligence is highly associated with the ability to rapidly model novel stimuli. Indeed, as Carpenter et al state in one early and prominent theoretical paper on intelligence: “analytic intelligence refers to the ability to deal with novelty” [63]. We could be measuring an indirect result of this modeling process by measuring the difference between the neural novelty response on the first viewing and later viewings. If the novelty response is no longer present in later viewings for a participant, presumably this is because this participant was able to successfully model this novel stimuli. Our adaptation measure could therefore be indirectly measuring how quickly or effectively participants model this novel movie scene.

Interestingly, one prominent neuroscientific theory of intelligence, the parieto-frontal integration theory of intelligence (P-FIT) [65–69] implicates a number of the same brain regions that were explored here. In particular, we find that central regions from this theory like the inferior and superior parietal lobule and the dorsal prefrontal cortex, which form the control system defined in [5, 70] form the backbone of the majority of our results, not only on intelligence and adaptation, but also on dimensionality/coupling changes and inter-subject synchronization, further suggesting a unifying factor underlying these results.

## Conclusion

In conclusion, our results suggest that a number of brain systems increase their connectivity and synchronize to incoming sensory stimuli in response to novelty. Additionally, we find evidence that this process is related to both psychological arousal and intelligence. These results can be seen in light of a broader emerging project to integrate intelligence and learning research into a network and systems perspective within neuroscience [14, 15, 18, 71–90]. In particular, we see these results as speaking to the participation of widely distributed brain systems in the recognition and accommodation to novel stimuli via model building.

## MATERIALS AND METHODS

### Human Connectome Project Data

The Human Connectome Project (HCP) 7T dataset [22] consists of structural magnetic resonance imaging (T1w), resting state functional magnetic resonance imaging (rsfMRI) data, movie watching functional magnetic resonance imaging (mwfMRI) from 184 adult subjects. These subjects are a subset of a larger cohort of approximately 1200 subjects additionally scanned at 3T. Subjects’ 7T fMRI data were collected during four scan sessions over the course of two or three days at the Center for Magnetic Resonance Research at the University of Minnesota. Subjects’ 3T T1w data were collected at Washington University in St. Louis. The study was approved by the Washington University Institutional Review Board and informed consent was obtained from all subjects.

### Demographics

We analyzed MRI data collected from *N*_*s*_ = 129 subjects (77 female, 52 male), after excluding subjects with poor quality data. Upon defining each spike as relative framewise displacement of at least 0.25 mm, we excluded subjects who fulfill at least 1 of the following criteria: greater than 15% of time points spike, average framewise displacement greater than 0.2 mm; contains any spikes larger than 5mm. Following this filter, subjects who contained all four scans were retained. At the time of their first scan, the average subject age was 29.36 ± 3.36 years, with a range from 22 − 36. 70 of these subjects were monozygotic twins, 57 were non-monozygotic twins, and 2 were not twins.

### MRI acquisition and processing

A comprehensive description of the imaging parameters and image preprocessing can be found in [91] and in HCP’s online documentation (https://www.humanconnectome.org/study/hcp-youngadult/document/1200-subjects-data-release). T1w were collected on a 3T Siemens Connectome Skyra scanner with a 32-channel head coil. Subjects underwent two T1-weighted structural scans, which were averaged for each subject (TR = 2400 ms, TE = 2.14 ms, flip angle = 8°, 0.7 mm isotropic voxel resolution). fMRI were collected on a 7T Siemens Magnetom scanner with a 32channel head coil. All 7T fMRI data was acquired with a gradient-echo planar imaging sequence (TR = 1000 ms, TE = 22.2 ms, flip angle = 45°, 1.6 mm isotropic voxel resolution, multi-band factor = 5, image acceleration factor = 2, partial Fourier sample = 7/8, echo spacing = 0.64 ms, bandwidth = 1924 Hz/Px). Four resting state data runs were collected, each lasting 15 minutes (frames = 900), with eyes open and instructions to fixate on a cross. Four movie watching data runs were collected, each lasting approximately 15 minutes (frames = 921, 918, 915, 901), with subjects passively viewing visual and audio presentations of movie scenes. Movies consisted of both freely available independent films covered by Creative Commons licensing and Hollywood movies prepared for analysis [92]. For both resting state and movie watching data, two runs were acquired with posterior-to-anterior phase encoding direction and two runs were acquired with anterior-to-posterior phase encoding direction.

Structural and functional images were minimally preprocessed according to the description provided in [91]. 7T fMRI images were downloaded after correction and reprocessing announced by the HCP consortium in April, 2018. Briefly, T1w images were aligned to MNI space before undergoing FreeSurfer’s (version 5.3) cortical reconstruction workflow. fMRI images were corrected for gradient distortion, susceptibility distortion, and motion, and then aligned to the corresponding T1w with one spline interpolation step. This volume was further corrected for intensity bias and normalized to a mean of 10000. This volume was then projected to the 2mm *32k fs LR* mesh, excluding outliers, and aligned to a common space using a multi-modal surface registration [93]. Resting state and moving watching fMRI images were nuisance regressed in the same manner. Each minimally preprocessed fMRI was linearly detrended, band-pass filtered (0.008-0.25 Hz), confound regressed and standardized using Nilearn’s signal.clean function, which removes confounds orthogonally to the temporal filters. The confound regression strategy included six motion estimates, mean signal from a white matter, cerebrospinal fluid, and whole brain mask, derivatives of these previous nine regressors, and squares of these 18 terms. Spike regressors were not applied. Following these preprocessing operations, the mean signal was taken at each time frame for each node, as defined by the Schaefer 200/400 parcellation(s) [5] in *32k fs LR* space.

### Fluid intelligence scores

Here we used fluid intelligence scores collected, processed and provided by the Human Connectome Project [22]. To access data, please see www.humanconnectome.org. Fluid intelligence was assessed using Penn Progressive Matrices, which is a shortened and computerized version of the Raven’s Progressive Matrices, a well-established measure of fluid intelligence that measures abstract reasoning using novel patterns in a pattern completion task [64].

### Quantifying dynamic psychological arousal

An independent sample of 46 undergraduate subjects watched the same complete set of movie-scenes as the Human Connectome Project cohort in order to provide a moment-by-moment rating of psychological arousal. These subjects were instructed to push a spring-loaded joystick (Pro Logitech Extreme 3D Pro; Model # 963290-0403) forward to indicate an increase in psychological arousal. A bar on the right side of the screen provided a real-time indicator of the degree to which they were moving the joystick, corresponding to a level between ‘not psychologically arousing’ (baseline; resting state of joystick) to ‘extremely psychologically arousing’ (pushed all the way forward). Subjects were instructed to continually rate their level of psychological arousal throughout each movie scene. These ratings were sampled at the presentation of each frame of the episode (24 samples per second). Because the repetition time (TR) of the fMRI scanner was 1 s, we used subjects’ mean arousal rating over each 1-s interval. Overall, this is similar to how other studies have used joysticks to get dynamic ratings of psychological responses to naturalistic stimuli (e.g. [94]). This study was approved by the institutional review board (IRB) at Indiana University.

## Acknowledgments

DPK was supported by NIMH R01MH110630.

---

**FIG. S1.**
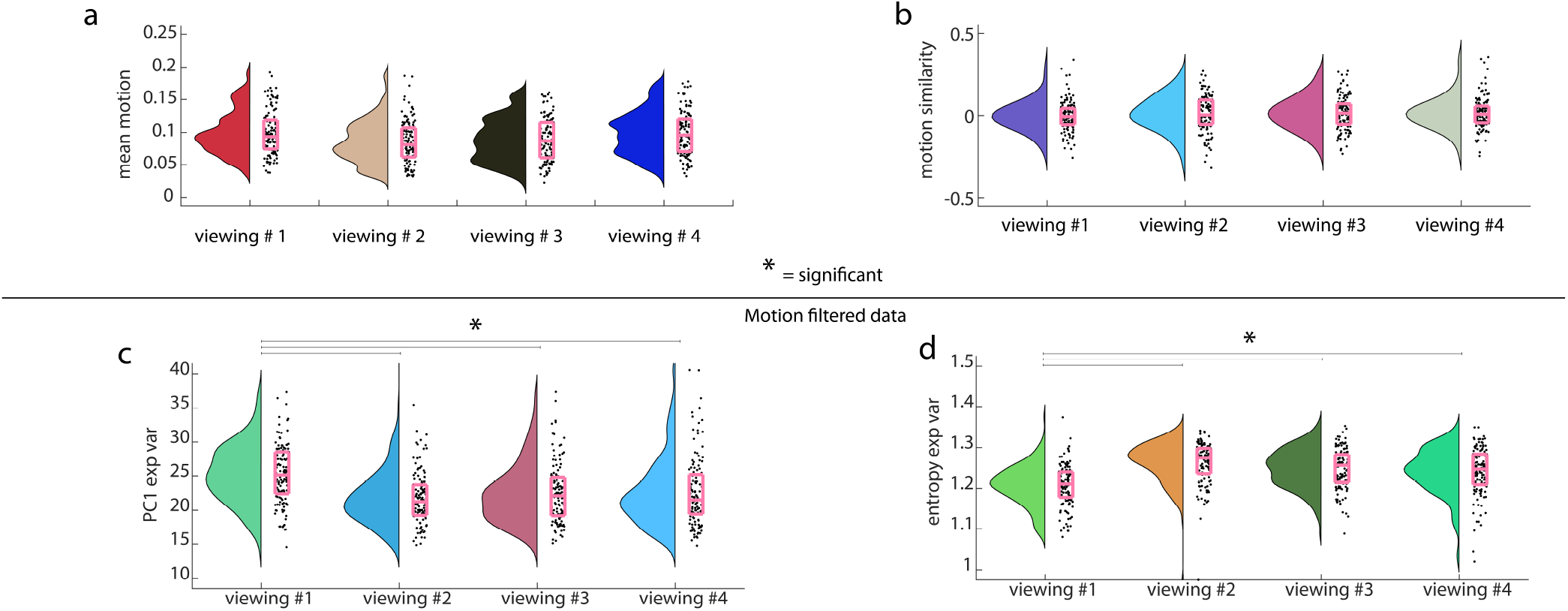
Novelty effect does not come from motion, (*a*) Box plots showing the mean motion per subject for each viewing. These distributions do not significantly differ. (*b*) Box plots showing the motion similarity across time per subject for each viewing. These distributions do not significantly differ. (*c*) Box plots showing that the explained variance of PC1 per viewing in motion filtered data (filter above 0.15 frame-wise displacement). First viewing distribution is significantly greater than later viewings (*p <* 0.05). (*d*) Box plots showing that the entropy of the explained variance per viewing in motion filtered data (filter above 0.15 frame-wise displacement). First viewing distribution is significantly lower than later viewings (*p <* 0.05).

**FIG. S2.**
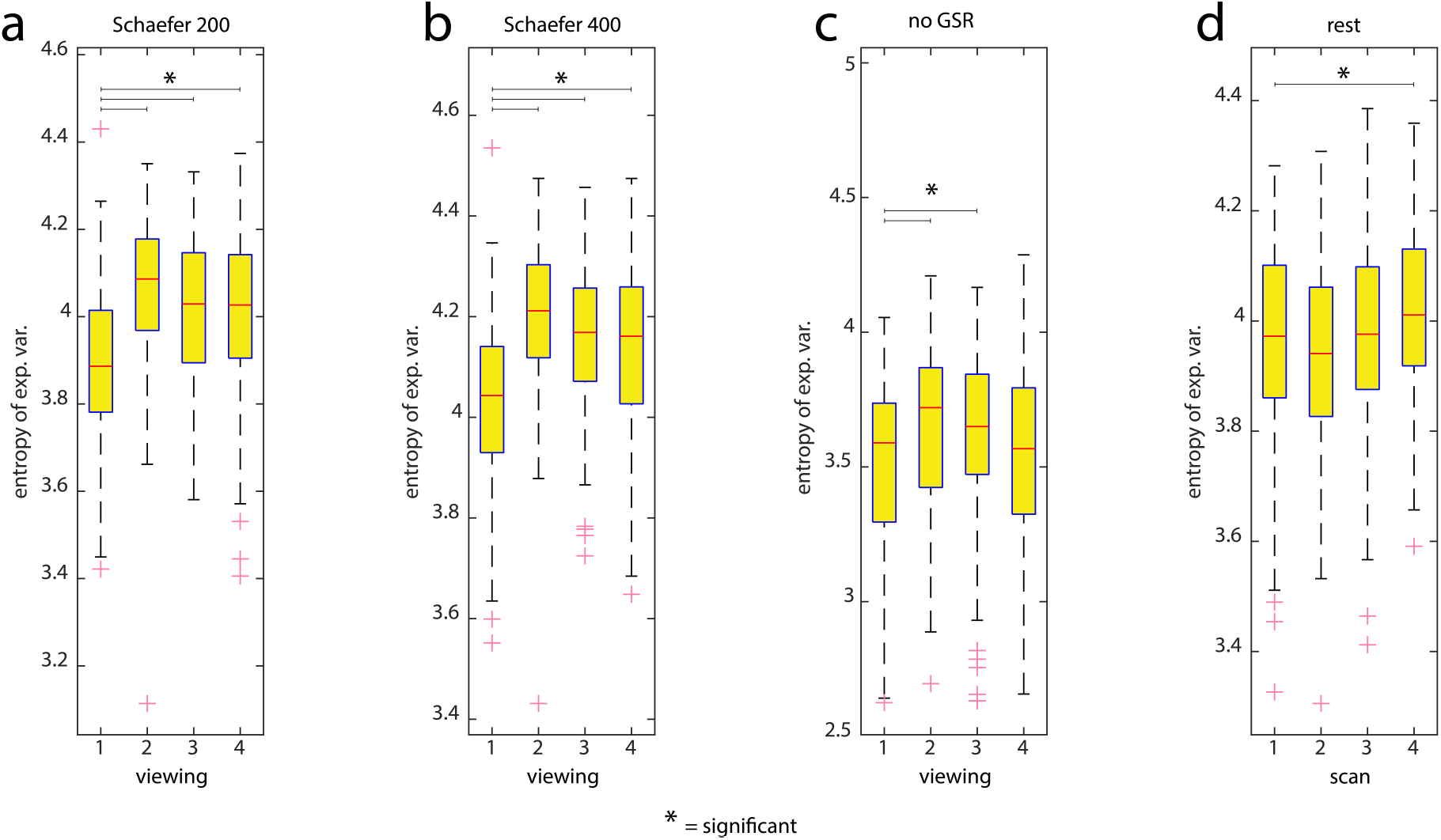
Novelty effect is found across scale and processing decisions, and not found at rest, (*a*) Box plots showing the entropy of the explained variance per viewing in Schaefer 200 parcellation (the parcellation we used in the main results). First viewing values are significantly lower than later viewings (*p <* 10^−5^). (*b*) Box plots showing the entropy of the explained variance per viewing in Schaefer 400 parcellation. First viewing values are significantly lower than later viewings (*p <* 10^−5^). (*c*) Box plots showing the entropy of the explained variance per viewing in data without global signal regression. First viewing values are significantly lower than the next two viewings, but not the final viewing (*p <* 0.05). (*d*) Box plots showing the entropy of the explained variance per scan in data from the same subjects during resting-state. First viewing values are significantly lower than later viewings (*p <* 0.05).

